# PRC1 resists microtubule sliding in two distinct resistive modes due to variations in the separation between overlapping microtubules

**DOI:** 10.1101/2024.12.31.630898

**Authors:** Daniel Steckhahn, Shane A. Fiorenza, Ellinor Tai, Scott Forth, Peter R. Kramer, Meredith Betterton

**Affiliations:** Department of Physics, University of Colorado Boulder, Boulder CO 80309, USA; Department of Physics, Faculty of Science, University of Zagreb, Bijenička Cesta 32, 10000 Zagreb, Croatia; Department of Biological Sciences and Center for Biotechnology and Interdisciplinary Studies, Rensselaer Polytechnic Institute, Troy NY 12180, USA; Department of Mathematical Sciences, Rensselaer Polytechnic Institute, Troy, NY 12180, USA; Department of Molecular, Cellular, and Developmental Biology, University of Colorado Boulder, Boulder CO 80309, USA

**Keywords:** microtubule, PRC1, kinesin, sliding, friction, geometry, resistance

## Abstract

Crosslinked cytoskeletal filament networks provide cells with a mechanism to regulate cellular mechanics and force transmission. An example in the microtubule cytoskeleton is mitotic spindle elongation. The three-dimensional geometry of these networks, including the overlap length and lateral microtubule spacing, likely controls how forces can be regulated, but how these parameters evolve during filament sliding is unknown. Recent evidence suggests that the crosslinker PRC1 can resist microtubule sliding by two distinct modes: a braking mode and a less resistive coasting mode. To explore how molecular-scale mechanisms influence network geometry in this system, we developed a computational model of sliding microtubule pairs crosslinked by PRC1 that reproduces the experimentally observed braking and coasting modes. Surprisingly, we found that the braking mode was associated with a substantially smaller lateral separation between the crosslinked microtubules than the coasting mode. This closer separation aligns the PRC1-mediated forces against sliding, increasing the resistive PRC1 force and dramatically reducing sliding speed. The model also finds an emergent similar average sliding speed due to PRC1 resistance, because higher initial sliding speed favors the transition to braking. Together, our results highlight the importance of the three-dimensional geometric relationships between crosslinkers and microtubules.

## INTRODUCTION

A hallmark of the cellular cytoskeleton is the generation of diverse architectures from a small number of building blocks. Polymer filaments, crosslinkers, motors, and associated proteins can self-organize into a range of assemblies that vary in size, mechanical stiffness, and dynamics. This allows dynamic cytoskeletal remodeling to facilitate intracellular transport, ciliary beating, chromosome segregation in mitosis, and cytokinesis, among other functions. A central question in cytoskeletal biology is how specific molecular interactions define the geometric organization of cytoskeletal assemblies and lead to emergent cytoskeletal dynamics and function.

A prototypical cytoskeletal assembly is the microtubule-based mitotic spindle, which is essential for successful chromosome segregation and cell division in eukaryotes. Structural motifs such as asters that emanate from spindle poles and antiparallel microtubule bundles that compose the interpolar spindle network are essential for providing the spindle with mechanical integrity, positioning chromosomes, and regulating spindle elongation. The relative positions and speeds of spindle microtubules are set by motor proteins that slide microtubules and non-motor proteins that crosslink and resist filament sliding. Important geometric parameters that regulate how forces are modulated include extent of overlap length, which is on the order of microns, and lateral distance between microtubule surfaces, which is typically tens of nanometers due to the typical length of crosslinking molecules. Coordination between the different spindle proteins regulate key aspects of spindle morphology, such as maintaining bipolarity and temporally regulating the overall length and width of the spindle.

Members of the MAP65 family of proteins, which includes Ase1 in yeasts, MAP65-1 in plants, and PRC1 in humans, preferentially crosslink microtubules in an antiparallel configuration. In dividing cells, they localize to the interpolar microtubule network beginning in metaphase and then become concentrated within overlaps throughout anaphase, acting as a brake against the motions of spindle pole separation and chromosome segregation(Pamula et al., 2019; Thomas et al., 2020; Mollinari et al., 2002; Loïodice et al., 2005; Jiang et al., 1998; Kajtez et al., 2016). These crosslinking proteins have also been shown to exhibit unique mechanical properties in reconstitution experiments. For example, Ase1 has been shown to act as an entropic spring to prevent complete sliding apart of antiparallel microtubules(Lansky et al., 2015), MAP65-1 can slow kinesin-driven microtubule gliding(Pringle et al., 2013), and PRC1 has been shown to act as a viscous dashpot by providing velocity-dependent frictional forces that oppose microtubule sliding(Gaska et al., 2020). Recent work from Alfieri et al. has suggested that PRC1 can adopt two distinct modes of resistive force production against kinesin-driving sliding forces(Alfieri et al., 2021). When molecules are evenly distributed throughout the overlap, PRC1 moderately slows filament sliding in a mode termed coasting. When PRC1 molecules become densely clustered, particularly near microtubule tips, a second mode termed braking abruptly emerges wherein the resistive forces increase and relative sliding velocity decreases significantly. In the braking mode, either PRC1-PRC1 or PRC1-microtubule interactions likely arise which are distinct from those during the coasting mode.

In addition to changes in overlap length along filaments during anaphase, variations in the lateral separation between crosslinked microtubules can also occur. Recent work found that the separation between pairs of motor-driven sliding microtubules changes with the sliding speed, suggesting that the angular tilt of motors within the overlap could affect sliding dynamics(Mitra et al., 2020; Meißner et al., 2024). Similarly, lateral microtubule separation has been measured by electron microscopy in the fission yeast mitotic spindle. This separation can be surprisingly small at 15 nm relative to the Ase1 crosslinker length of 32 nm, suggesting that crosslinkers can be compacted or tilted within overlaps(Ding et al., 1993; Ward et al., 2015). However, the binding geometry and orientation of PRC1 during microtubule pair sliding has not been directly measured.

Here, we sought to understand what molecular mechanisms contribute to PRC1 braking and its enhanced resistive force production during microtubule sliding, how microtubule pairs can rapidly transition from coasting to braking, and how the geometry between filaments might be regulated during this process. Stochastic modeling of protein binding to and movement on microtubules can help elucidate these underlying mechanisms. Previous work has modeled friction of proteins bound to microtubules, crosslinkers on microtubule pairs, and formation of microtubule overlaps (Bormuth et al., 2009; Forth et al., 2014; Lansky et al., 2015; Johann et al., 2015, 2016; Kuan and Betterton, 2016; Prelogović et al., 2019; Wijeratne and Subramanian, 2018; Lera-Ramirez and Nedelec, 2019; Winters et al., 2019; Lamson et al., 2019; Krattenmacher et al., 2024). We used the Cytoskeleton Lattice-based Kinetic Simulator (CyLaKS), designed to study systems of microtubules and crosslinkers where details such as crosslinker interactions and microtubule separation are important (Fiorenza et al., 2021; Wijeratne et al., 2022). We previously found that crosslinkers modulate the separation between crosslinked microtubule pairs (Fiorenza et al., 2021). Here, we model resistive force from PRC1 during motor-driven microtubule sliding. We find that two geometrically distinct modes of sliding occur, with different typical crosslinker tilt in the overlap and microtubule separation. Increased crosslinker angle and closer separation coincide with the transition from coasting to braking, an effect which is more pronounced when unloaded microtubule sliding velocities are higher. Our results suggest that lateral microtubule separation is an under-appreciated determinant of force in crosslinked microtubule bundles.

## RESULTS

### Computational modeling reproduces two modes of PRC1 resistance in motor-driven microtubule overlaps

Recent work examined how microtubule pairs crosslinked by PRC1 responded to surface-bound kinesin-1 sliding the microtubules apart (Alfieri et al., 2021) (Fig. 1A-D). During coasting events, PRC1 was approximately uniformly distributed within the overlap (Fig. 1B), and the sliding speed was only moderately reduced from free microtubule sliding (Alfieri et al., 2021). During braking events, PRC1 clustered near overlap edges (Fig. 1C) and strongly resisted sliding. The effect of PRC1 sliding resistance can be quantified by comparing the average sliding velocity for overlapping microtubules, V_bundled_, to the average sliding velocity after the microtubules were no longer crosslinked by PRC1, V_escaped_. This normalized velocity V_bundled_*/*V_escaped_ showed a bimodal distribution with the low velocity ratio group representing braking and the high speed group coasting (Fig. 1D) (Alfieri et al., 2021).

**Figure 1:**
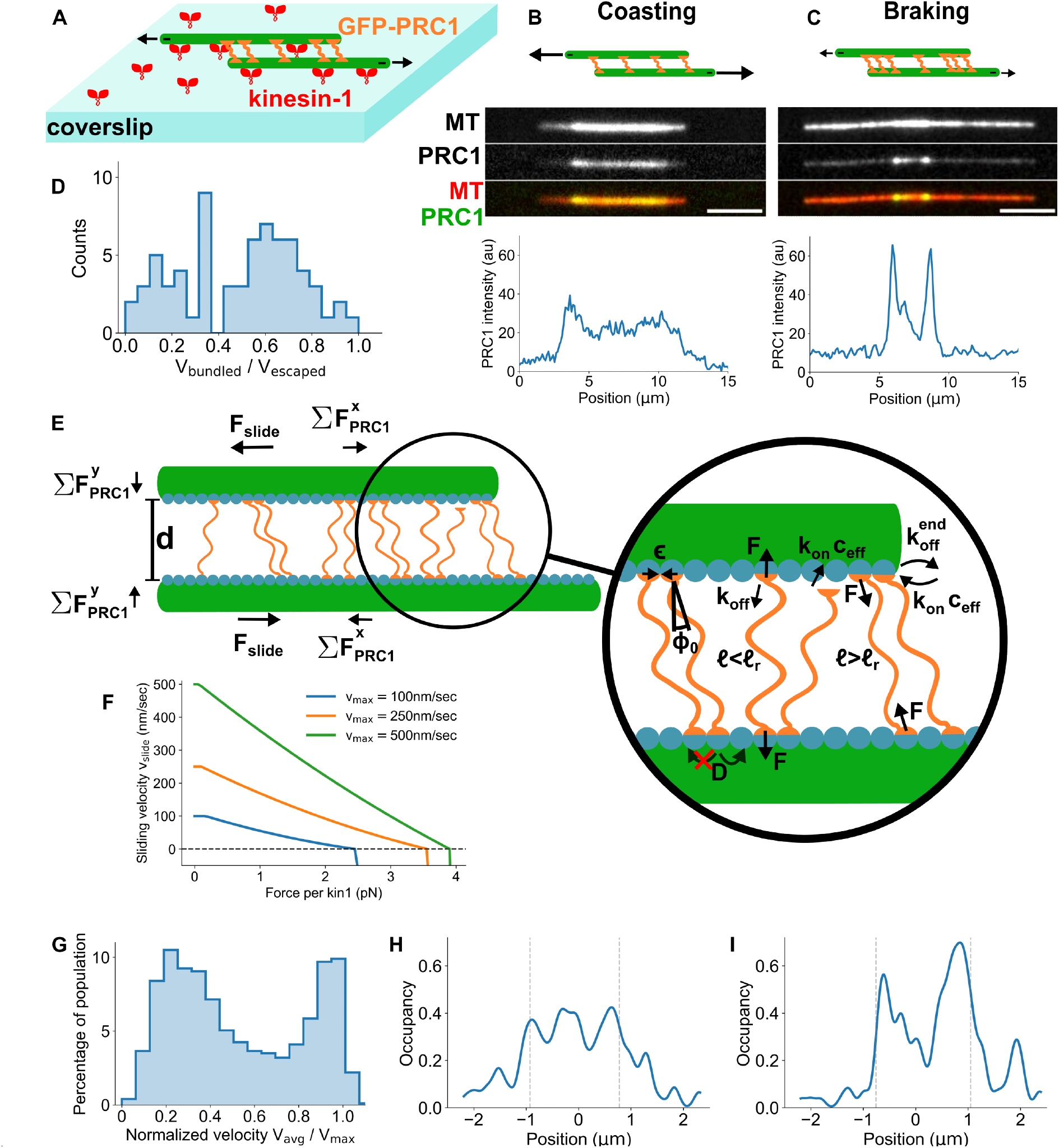
A computational model of PRC1 resistance to microtubule overlap sliding reproduces experimentally observed braking and coasting modes. (A) Schematic of experimental microtubule pair sliding of Alfieri et al. (2021) Anti-parallel microtubules crosslinked by PRC1 were slid apart by surface-attached kinesin-1. (B-C) Schematic (top), TIRF images (middle), and fluorescence intensity line scans (bottom) of two representative microtubule overlaps showing coasting (B) and braking (C) events. (D) Distribution of the normalized velocity V_bundled_*/*V_escaped_ for microtubule pairs and an ATP concentration of 10 *μ*M. (E) Schematic of computational model geometry and ingredients. (F) Microtubule sliding velocity as a function of PRC1 force per kinesin motor used in our sliding model for different values of the unloaded kinesin velocity. The horizontal line at v_slide_ = 0 shows where the velocity model switches to the backsliding model. (G) Distribution of the normalized velocity V_avg_*/*V_max_ from simulations with a distribution of maximum filament velocity corresponding to experiments with 10 *μ*M ATP. (H,I) PRC1 occupancy in simulated overlaps (edges marked by dashed lines) for the coasting (H) and braking (I) state.

To explore the molecular mechanisms responsible for the two resistive modes of PRC1 we developed a computational model of microtubule sliding resisted by crosslinking PRC1 molecules (Methods, Fig. 1E). In our model, overlapping microtubules crosslinked by PRC1 move due to force from PRC1, kinesin-driven sliding, steric repulsion, and random thermal kicks. Microtubules move both longitudinally (*x* direction in Fig. 1E, along their length) and laterally (*y* direction in Fig. 1E, across the microtubule separation).

PRC1 molecules can bind to, unbind from, and diffuse between discrete sites along the microtubules. When crosslinking, PRC1 molecules exert linear spring-like forces on the microtubules in response to stretching and compression. In addition, previous experimental work revealed a tilt of the PRC1 molecular axis relative to the microtubule (Subramanian et al., 2010; Kellogg et al., 2016) and AlphaFold modeling also suggests a relative tilt (Fig. S1). Therefore, we modeled PRC1 with a preferred tilt angle and a spring-like restoring torque for deviations away from the preferred angle (Methods). In what follows, we refer to the force along the molecular axis as the “linear force” and the force induced on the heads by the restoration toward the preferred tilt angle as the “torsional force.” PRC1 diffusion, binding, and unbinding are force dependent consistent with Boltzmann statistics. PRC1 molecules interact with each other through steric interactions that prevent PRC1 heads from occupying the same site or crossing. PRC1 also have a neighbor-neighbor (N-N) attractive interaction between neighboring PRC1 heads. PRC1 heads are restricted from diffusing off microtubule ends, but end-bound PRC1 heads have an increased rate of unbinding. Kinesin-driven sliding is modeled via an effective relationship between the resistive force from the PRC1 and the velocity at which the microtubule is driven by surface bound kinesin-1 (Methods, Fig. 1F). We initialized simulations with two anti-parallel microtubules crosslinked by PRC1. After an initial binding equilibration period during which sliding is turned off, we allowed sliding and ran simulations until the microtubules slid apart.

Our model reproduced the braking and coasting modes observed experimentally, with a bimodal velocity distribution and variable PRC1 intensity at overlap edges (Fig. 1G-I). We used the normalized velocity V_norm_ = V_avg_*/*V_max_ to determine the velocity distribution, where V_avg_ is the average speed during overlap sliding and V_max_ is the kinesin-driven sliding speed of a free microtubule (Fig. 1G). We then used the model to further investigate the mechanisms that distinguish coasting and braking.

### PRC1 accumulates on overlap edges at the onset of braking

Since PRC1 accumulation on the edge of overlaps was correlated with braking in experiments (Alfieri et al., 2021), we analyzed the temporal dynamics of PRC1 overlap concentration and sliding speed in our model. During coasting events the sliding speed remained near V_max_ and PRC1 was not consistently concentrated near overlap edges (Fig. 2A). In our simulations overlaps transitioned abruptly from coasting to braking, apparent as the significant decrease in the velocity. This sliding velocity decrease was accompanied by PRC1 accumulation at overlap edges (Fig. 2B, S2). This demonstrates that the formation of dense PRC1 clusters at overlap edges temporally coincided with a decrease in sliding speed in our model, consistent with experimental observations (Alfieri et al., 2021). The accumulation of PRC1 at overlap edges starts with overlap shortening that sweeps crosslinkers closer to the edges (Fig. 2C). PRC1 neighbor-neighbor interactions tend to maintain clusters, while the unbinding of PRC1 heads or their diffusion away from edges decreases clustering (Fig. 2D). The balance between sliding, neighbor attraction, diffusion, and unbinding determines whether edge clusters of PRC1 form and are maintained.

**Figure 2:**
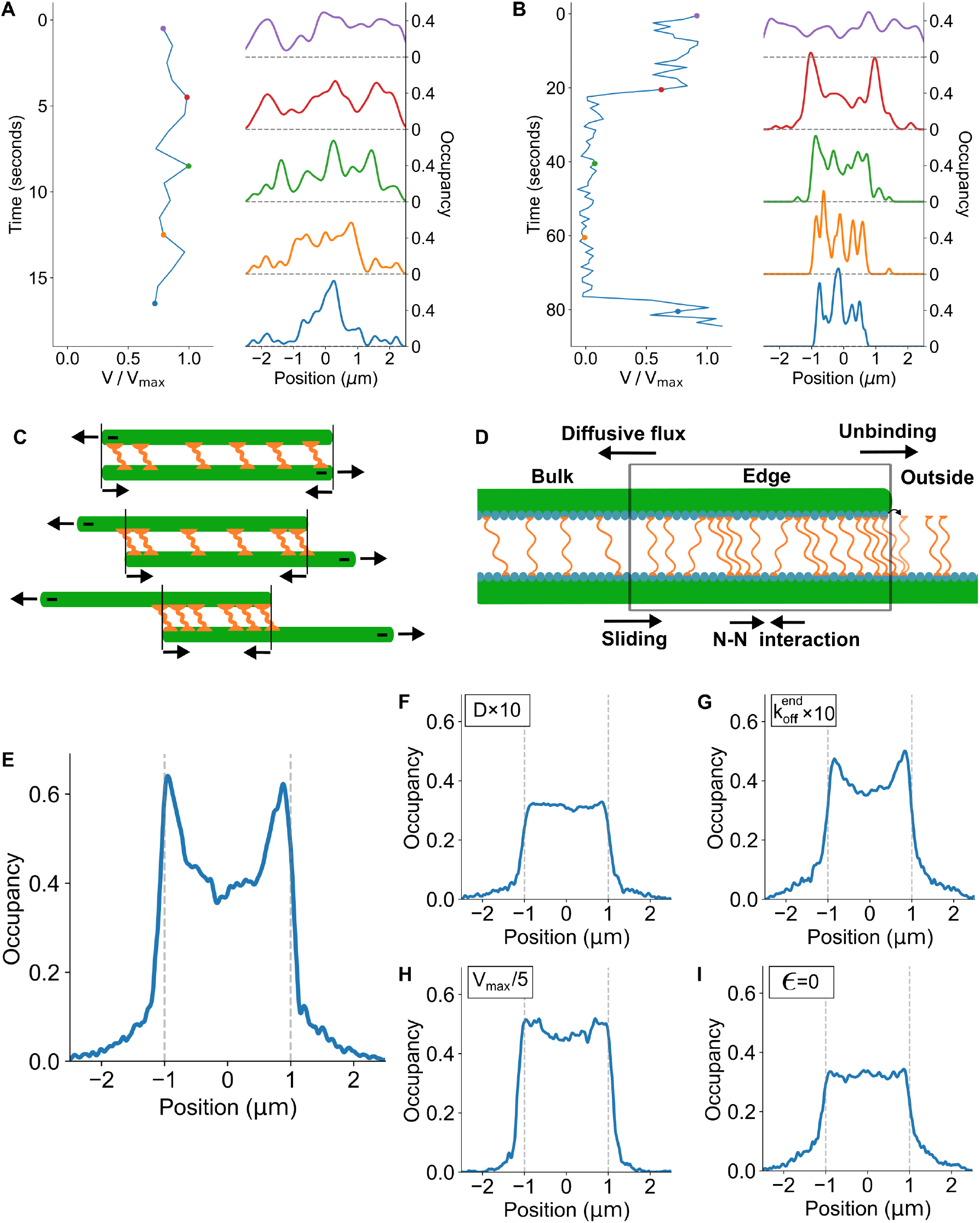
The transition from coasting to braking is associated with PRC1 movement to overlap edges and clustering. (A-B) Left, instantaneous normalized velocity V(t)*/*V_max_, for simulations showing coasting (A) and braking (B). Right, average PRC1 lattice occupancies shown at the times indicated by colored points on the velocity-time curve. (C) Schematic of PRC1 accumulation at overlap edges. (D) Schematic of molecular mechanisms that contribute to PRC1 movement into and out of the overlap edge. (E-I) Average PRC1 lattice occupancy for 2 *μ*m-long simulated overlaps, from 100 simulations at the time when the overlap length first decreased below 2 *μ*m. Results for the reference simulation parameter set of Tables 1, 2 and V_max_=50 nm/sec (E) and simulations with one parameter varied (F-I). (F) Diffusion coefficient of crosslinking PRC1 increased by a factor of 10 to 0.4 *μ*m s^*−*1^ (G) End unbinding rate increased by a factor of 10 to 1 s^*−*1^. (H) Sliding speed decreased by a factor of 5 to 10 nm/sec. (I) Neighbor-neighbor interaction energy between PRC1 molecules decreased to 0.

To test this conceptual picture, we varied model parameters that control movement into/out of the overlap edges (Fig. 2E-I). Increasing the diffusion coefficient or end unbinding rate both tend to remove PRC1 movement from overlap edges, and in simulations increasing either parameter decreased edge cluster formation (Fig. 2F,G). Reducing the kinesin-driven sliding velocity or turning off the N-N interactions tend to lower the rate of PRC1 accumulation or persistence at overlap edges. As expected, these parameter changes also decreased edge clustering in our simulations (Fig. 2H,I). These results support our picture of the factors that control PRC1 edge clustering (Fig. 2D).

**Table 1:** Simulation parameters for microtubules and PRC1.

| Quantity | Symbol | Value | Notes |
| --- | --- | --- | --- |
| Time step | $\Delta t$ | 0.00002 sec | Simulation stability; resolution of binding kinetics |
| Maximum simulation time | $t_f$ | 300 sec | |
| Data collection time | $\Delta t_c$ | 1 sec | |
| Equilibration time | $\Delta t_i$ | 1.5 sec | |
| Thermal energy | $k_B T$ | 4.1 pN nm | Room temperature |
| **Microtubules** |  |  |  |
| Radius | R | 12.5 nm | Howard (2001) |
| Length | $L_m$ | 5 $\mu\text{m}$ | Alfieri et al. (2021) |
| Site spacing | $\delta$ | 8 nm | Howard (2001) |
| Longitudinal diffusion coefficient | $D_{\parallel}$ | $V_{\max} \times 0.18 \text{ nm}$ | See text |
| Lateral diffusion coefficient | $D_{\perp}$ | $V_{\max} \times 0.09 \text{ nm}$ | See text |
| Longitudinal drag coefficient | $\gamma_{\parallel}$ | $V_{\max}^{-1} \times 23 \text{ pN sec nm}^{-1}$ | See text |
| Lateral drag coefficient | $\gamma_{\perp}$ | $V_{\max}^{-1} \times 46 \text{ pN sec nm}^{-1}$ | See text |
| WCA potential strength | $\epsilon_{\text{WCA}}$ | 5 pN nm | See text |
| WCA interaction scale | $\sigma$ | 45 nm | See text |
| **PRC1** |  |  |  |
| Average number | N | 100 | Alfieri et al. (2021) |
| Singly bound diffusion coefficient | $D_s$ | $0.13 \mu\text{m}^2 \text{ s}^{-1}$ | van den Wildenberg (2011) |
| Doubly bound diffusion coefficient | D | $0.04 \mu\text{m}^2 \text{ s}^{-1}$ | van den Wildenberg (2011), Fiorenza et al. (2021) |
| Neighbor-neighbor energy | $\epsilon$ | 2.3 $k_B T$ | Estimated from van den Wildenberg (2011) |
| Spring constant | $k_s$ | 2 pN/nm | Fit, see text |
| Rest length | $\ell_o$ | 32 nm | Subramanian et al. (2013) |
| Binding rate | $k_{\text{on}}^s$ | $0.000238 \text{ nM}^{-1} \text{ s}^{-1}$ | Subramanian et al. (2010) |
| Bulk concentration | $c_{\text{bulk}}$ | 1 nM | Alfieri et al. (2021) |
| Binding rate of 2<sup>nd</sup> head | $k_{\text{on},2}^s$ | $5.88 \text{ s}^{-1}$ | Fiorenza et al. (2021) |
| Singly bound unbinding rate | $k_{\text{off}}^s$ | $0.1 \text{ s}^{-1}$ | Subramanian et al. (2010) |
| Doubly bound unbinding rate | $k_{\text{off}}$ | $0.1 \text{ s}^{-1}$ | Subramanian et al. (2010) |
| Unbinding rate at microtubule ends | $k_{\text{off}}^{\text{end}}$ | $20 \text{ s}^{-1}$ | Fit, Fig. S8 |
| Preferred angle | $\phi_0$ | 20° | Subramanian et al. (2010), Fig. S1 |
| Angular spring constant $-\phi$ | $k_{\phi-}$ | $45 \text{ pN nm rad}^{-1}$ | Fit, Fig. S4 |
| Angular spring constant $+\phi$ | $k_{\phi+}$ | $3 \text{ pN nm rad}^{-1}$ | Fit, Fig. S5 |

**Table 2:** Model parameters for kinesin.

| Quantity | Symbol | Value | Notes |
| --- | --- | --- | --- |
| **Kinesin** |  |  |  |
| Number of available kinesin | $N_{\text{kin}}$ | 30 | see text |
| Unloaded microtubule gliding velocity | $V_{\text{max}}$ | 100-500 nm s<sup>-1</sup> | Schnitzer et al. (2000); Alfieri et al. (2021) and text |
| Stall force | $F_s$ | 6 pN | Ma et al. (2023) |
| Step size | $\delta$ | 8 nm | Carter and Cross (2005) |
| Tether compliance | $\kappa$ | 0.25 pN nm<sup>-1</sup> | Furuta et al. (2013) |
| Rest length | $\ell_0$ | 25 nm | see text |
| Reattachment rate | $k_a$ | 5 s<sup>-1</sup> | Klumpp et al. (2015) |
| Detachment rate under low hindering load | $k_{d+}^0$ | 1.11 sec<sup>-1</sup> | Andreasson et al. (2015a) |
| Detachment rate under assisting load | $k_{d-}^0$ | 7.4 sec<sup>-1</sup> | Andreasson et al. (2015a) |
| Force scale of accelerated detachment under hindering load | $F_{d+}$ | 12 pN | Andreasson et al. (2015a) |
| Force scale of accelerated detachment under assisting load | $F_{d-}$ | 22 pN | Andreasson et al. (2015a) |

### Braking and coasting overlaps show differences in PRC1 tilt and microtubule separation

Beyond PRC1 clustering at overlap edges, we asked whether braking and coasting overlaps show other differences. Changes in microtubule lateral separation alter the compression/extension and angle of PRC1, potentially altering both the magnitude and direction of PRC1 force (Fig. 3A). Analysis of two microtubules in a simulated overlap with a braking event showed a transition from ∼30 nm separation between the surfaces of the two microtubules to ∼15 nm at the same time that the sliding velocity dropped (Fig. 3B). We then examined sliding velocity, microtubule separation, and PRC1 angle for multiple simulations and found different overlap and PRC1 geometry in the braking and coasting states (Fig. 3C-E). As shown in these diagrams, we measure the tilt angle relative to the normal direction between the microtubules, with positive values corresponding to the tilt naturally induced by the sliding. In the braking state, sliding velocity was low, microtubule separation was ∼10-15 nm, and the PRC1 tilt angle was ∼64.8 degrees (Fig. 3C,D). In the coasting state, sliding velocity was high, microtubule separation was ∼32 nm (the length of PRC1), and the mean PRC1 tilt angle was ∼13.6 degrees (Fig. 3D, E). To further test whether lower microtubule separation caused lower sliding speed, we simulated overlaps with fixed separation at the average values for braking (14.0 nm) and coasting (30.8 nm) overlaps. The resulting overlap sliding speed for fixed-separation experiments were comparable to those found in our variable-separation braking and coasting states. (Fig. 3D, S3). This shows that in our model, the reduced separation of the braking state causes the reduction in sliding speed.

**Figure 3:**
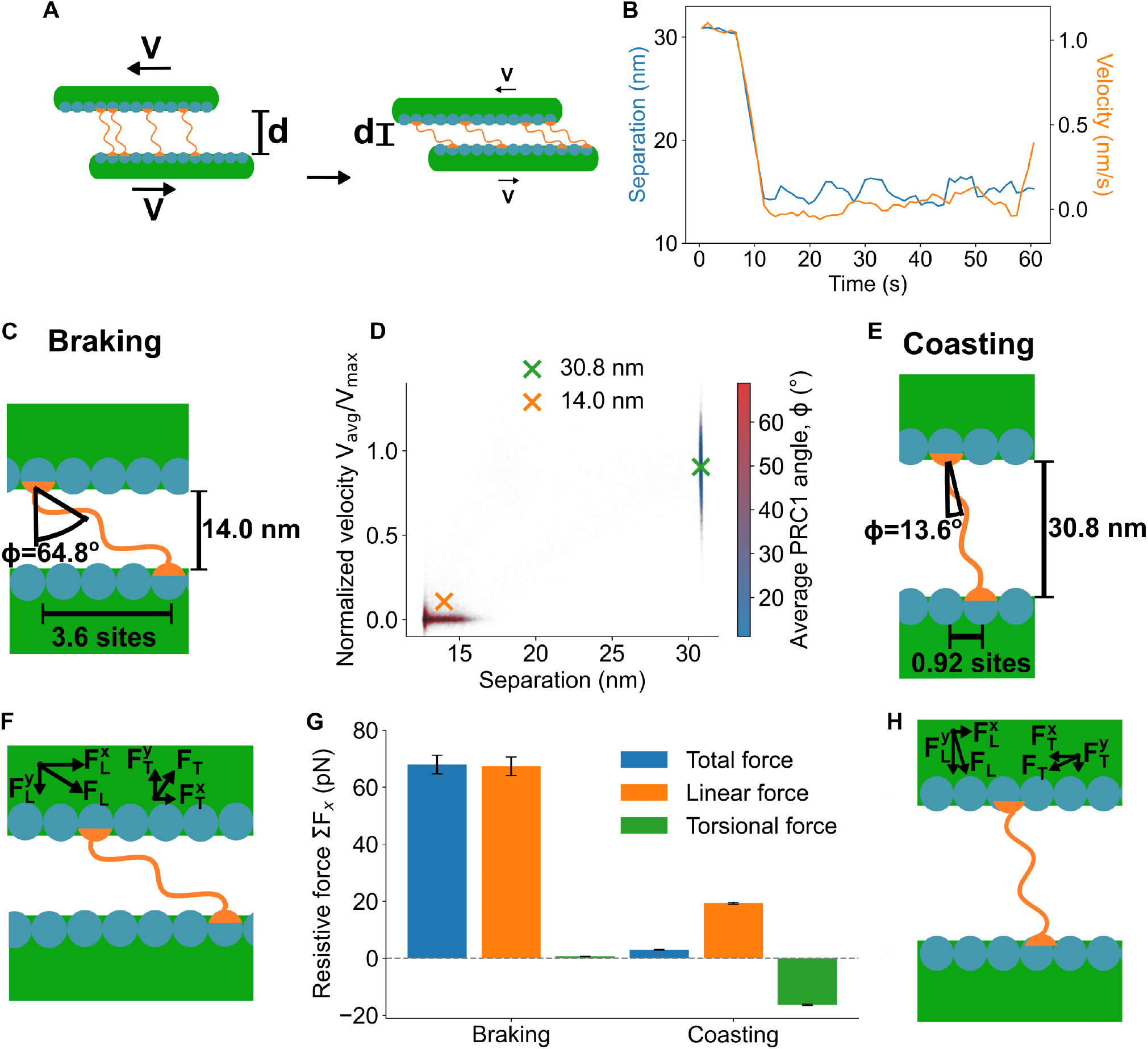
Braking and coasting states show distinct overlap geometry with different lateral microtubule separation and crosslinker tilt. (A) Schematic of coasting (left) and braking (right) states, showing decreased microtubule separation *d* and increased crosslinker tilt in the braking state. (B) Microtubule lateral separation (left axis, blue) and overlap sliding speed (right axis, orange) as a function of time when a braking event occurs. Sliding speed and microtubule separation decrease at the same time. (C) Schematic of microtubule and crosslinker geometry in the braking state. The average microtubule separation is 14 nm, PRC1 tilt angle is on average 64.8^*°*^, and PRC1 heads are displaced longitudinally along the microtubule by an average of 3.6 sites. (D) Normalized sliding velocity (left axis) and PRC1 angle (color) as a function of microtubule pair separation. Points show snapshots of unconstrained simulations, green and orange crosses show average speeds across simulations with fixed microtubule lateral separation. (E) Schematic of microtubule and crosslinker geometry in the coasting state. The average microtubule separation is 30.8 nm, PRC1 tilt angle is on average 13.6^*°*^, and PRC1 heads are displaced longitudinally along the microtubule by an average of 0.92 sites. (F) Schematic of microtubule and crosslinker geometry in the braking state showing the typical direction and magnitude of the linear spring force *F*_*L*_ and torsional spring force *F*_*T*_. (G) Average total PRC1 resistive force, and its torsional and linear components in the braking and coasting states. (H) Schematic of microtubule and crosslinker geometry in the coasting state showing the typical direction and magnitude of the linear spring force *F*_*L*_ and torsional spring force *F*_*T*_.

The linear and torsional contributions to the PRC1 force change with the angle of PRC1 (Fig. 3F-H). The braking state has more steeply tilted PRC1 that align the linear force of PRC1 extension (or compression) in the longitudinal direction, while the longitudinal component of torsional force is relatively small (Fig. 3F, G). By contrast, in the coasting state most PRC1 molecules are less tilted than the preferred angle (Fig. 3G, H, S4A,B). As a result the linear force along the axis of PRC1 molecules is primarily lateral while the torsional force perpendicular to the molecule axis is primarily longitudinal. The longitudinal components of the linear and torsional forces are opposed, largely canceling each other and leading to a small net resistive force. As a result, the change in PRC1 angle between coasting and braking leads to a factor of ∼20 change in the resistive force (Fig. 3G).

### The clustering of PRC1 at overlap edges pulls microtubules closer together

We often observed in simulations that clustering of PRC1 at overlap edges preceded the decrease in microtubule separation and transition to braking (Fig. 2B, 4A). Our results can be explained by differences in PRC1 molecules oriented with sliding (+*ϕ*, Fig. 4B) and PRC1 molecules oriented against sliding (-*ϕ*). Antiparallel microtubule sliding in the overlap tends to rotate PRC1 molecules toward more positive values of *ϕ*, orienting them with sliding (Fig. 4B). PRC1 molecules oriented with sliding tend to be stretched by sliding and will therefore exert a lateral force that pulls microtubules together. The lateral force arising from PRC1 torsion is negligible for molecules oriented with sliding (Fig. S5). Conversely, molecules oriented against sliding tend to be compressed by sliding, and will tend to exert a lateral force that pushes microtubules apart, due to contributions from both the linear and torsional components of force(Fig. 4B, S5).

**Figure 4:**
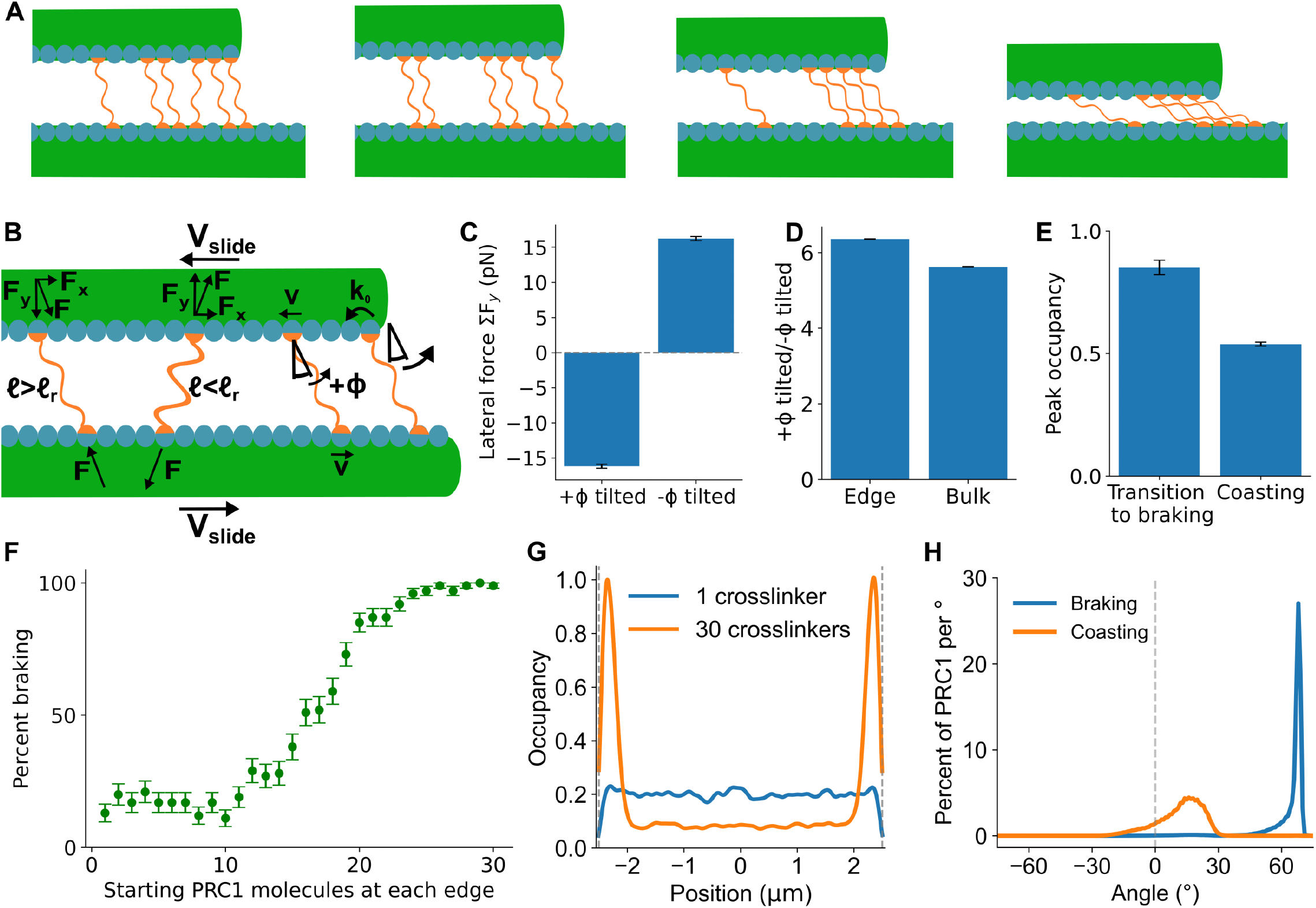
PRC1 accumulation at overlap edges causes the transition from coasting to braking. (A) Example time-lapse from a simulation showing the orientation of edge PRC1 molecules as an overlap transitions from braking to coasting (0.02 seconds between frames). (B) Schematic of length, tilt angle, and force produced by PRC1 tilted in the direction of sliding (labeled +*ϕ*) and against sliding (*−ϕ*). Sliding typically causes PRC1 tilted with sliding to extend and exert force that decreases microtubule separation, while PRC1 tilted against sliding typically becomes compressed and exert force that increases microtubule separation. (C) Average lateral PRC1 force from subpopulations of PRC1 molecules tilted with and against sliding, from simulations in the coasting state. (D) Ratio of the number of PRC1 molecules tilted with and against sliding for edge and bulk regions. (E) Maximum PRC1 occupancy at overlap edges at the time when overlaps transitioned from coasting to braking and during coasting. In B-D data are averages from 1000 simulations. (F) Percentage of simulations that transition to braking as a function of the number of PRC1 molecules initially present at overlap edges. Points show mean and standard error from 100 simulations. (G) Average initial PRC1 occupancy for simulations with 0 edge PRC1 and 30 edge PRC1. (H) Angular distribution of PRC1 molecules during braking and coasting.

To test this intuition, we measured the PRC1 lateral force exerted by molecules of different tilt direction in our simulations, and found a net negative (attractive) lateral force for PRC1 molecules oriented with sliding and a net positive (repulsive) lateral force for PRC1 molecules oriented against sliding (Fig. 4C). Second, PRC1 molecules on the overlap edges preferentially orient with sliding, since a PRC1 molecule head at an overlap edge can only hop in a direction that increases the +*ϕ* tilt (Fig. 4B). As PRC1 accumulates near overlap edges, steric exclusion will prevent PRC1 from diffusing toward the microtubule plus ends, further contributing to positive tilt. Consistent with this, the ratio of positive to negative tilt angle is higher for PRC1 molecules near overlap edges (Fig. 4D). Therefore, as PRC1 accumulates at edges, the number of PRC1 with positive tilt increases, which creates forces that pull the microtubules together and drive a transition to lower sliding speed. As the overlaps transition to braking the PRC1 molecules are driven to angles greater than their preferred angle, indicating that the linear attractive force is primarily responsible for the transition to braking.

This picture predicts that PRC1 will be concentrated at overlap edges just before the speed decrease from coasting to braking. We measured the maximum of smoothed PRC1 occupancy in the edge region, and found that more PRC1 was present at the overlap edges during the transition from coasting to braking than on average during coasting (Fig. 4E). To confirm that the buildup of PRC1 on overlap edges causes transitions to the braking state and is not just correlated with it, we ran simulations where overlaps started with a varying amount of PRC1 at the overlap edge regions, while keeping the total number of PRC1 molecules constant. The fraction of simulations that transitioned from the braking to coasting state within the first 10 seconds of the simulation increased with the initial number of edge PRC1 molecules (Fig. 4F-G), indicating that PRC1 edge accumulation drives the transition to braking.

In addition to explaining why overlaps transition from coasting to braking, the difference in behavior between crosslinkers tilted with and against sliding explains why overlaps remain in the braking state. In coasting overlaps, crosslinkers are tilted both with and against sliding, causing a balance of forces (Fig. 4B, H). In contrast, in the braking state crosslinkers are only tilted with sliding, pulling the microtubules together (Fig. 4 B,H). This is due to the high energy required for crosslinkers to be tilted against sliding (Fig. S5).

### PRC1 resistance to sliding narrows the distribution of overlap sliding speed

Experiments on sliding of overlaps with PRC1 found that higher sliding speed (due to increased ATP concentration) led to more frequent braking events (Alfieri et al., 2021). To test whether this occurs in our model, we simulated sliding overlaps with a range of maximum sliding speed of 50-500 nm/sec (Fig. 5A). Simulated overlaps with higher maximum sliding speed showed a greater decrease in speed due to PRC1 resistive force, consistent with experimental observations. To further test this effect, we ran simulations with maximum velocity were sampled from Gaussian fits of the experimental velocity distributions with 10 *μ*M, 100 *μ*M, and 500 *μ*M ATP (Fig. 5B) (Alfieri et al., 2021). For faster sliding speed, overlap sliding decreased more (Fig. 5C). This occurs because overlaps with faster sliding are more likely to accumulate PRC1 molecules at overlap edges (Fig. 2H) and are therefore more likely to switch from coasting to braking. This effect narrowed the distribution of overlap sliding speed. The average maximum sliding velocity varied by a factor of ∼3 for the simulated distributions corresponding to 10-500 *μ*M ATP concentration (means of 127, 257, and 406 nm/s, Fig. 5B), while the overlap sliding speeds varied by only a factor of ∼2 (means of 65, 58, and 36 nm/sec, respectively, Fig. 5C). This suggests that the ability of PRC1 to resist sliding through both coasting and braking reduces the variability in overlap sliding speed, helping regulate sliding velocity to a narrower range. This is consistent with experimental results showing that the sliding velocity has a weaker dependence on ATP concentration than the single-microtubule velocity. (Alfieri et al., 2021)

**Figure 5:**
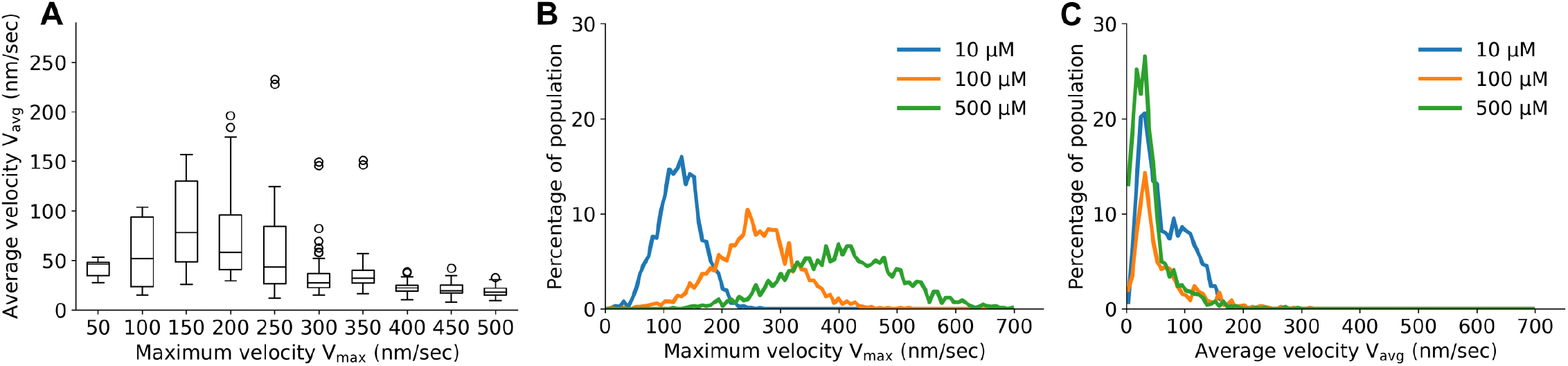
The two resistive modes of PRC1 lead to similar overlap sliding speeds for a range of single-microtubule sliding speed. (A) Average overlap sliding velocity as a function of single-microtubule sliding velocity. Data shown from 100 simulations for each condition. (B) Histogram of single-microtubule maximum velocity adapted from experimental results of Alfieri et al. (2021) for varying ATP concentration. (C) Histogram of overlap sliding velocity corresponding to the same simulations show in (B). Data shown from 1000 simulations.

## DISCUSSION

We used computational modeling to study how crosslinking PRC1 molecules resist sliding of antiparallel microtubule overlaps. Consistent with experimental work (Alfieri et al., 2021), our model showed that crosslinker-driven resistance to sliding occurred in two distinct modes: a coasting state with a slight speed decrease and a braking state with a larger slowdown. Our computational approach allowed a dissection of the mechanisms that distinguished the coasting and braking states, as well as a description for how transitions between the two modes might occur.

We found that the slower braking state was characterized by accumulation of PRC1 at the overlap edges, which tended to drive PRC1 to higher tilt angle and concomitantly smaller lateral separation between microtubule surfaces. When the microtubules were closer together, the crosslinkers exert a higher longitudinal force to resist sliding. Remarkably, our results suggest that the braking and coasting states are structurally and geometrically distinct. The coasting state (with microtubule sliding velocities more than 40% of the uncrosslinked values) has microtubules separated by *>*30 nm and PRC1 angle relative to the normal to the microtubule of *<*20 degrees, which is consistent with PRC1 orientation observed in cryoEM tomography studies (Subramanian et al., 2010). The transition to braking is characterized by an abrupt transition to slower speed (with sliding velocity less than 40% of the uncrosslinked value), microtubule separation ∼15 nm, and PRC1 angle *>*50 degrees. This sharp change reflects a dramatic rearrangement of the crosslinkers and resembles a phase transition.

Our work extends and helps explain the literature linking microtubule separation and force exerted by crosslinking motors and MAPs. Neighboring microtubules in the fission-yeast late anaphase mitotic spindle midzone are separated by ∼15 nm (Ding et al., 1993; Ward et al., 2015), lower than in early mitosis. The functional significance of this spacing decrease to at least a factor of two smaller than the length of the midzone crosslinkers and motors has remained unclear. Our results suggest a possible explanation: the resistance of Ase1 crosslinkers to spindle microtubule sliding as the spindle elongates could drive a transition to a braking-like low separation/high tilt state. The 15 nm microtubule separation is similar to what we predict for microtubule pairs that have engaged in a braking mode. While Ase1 has been shown to generate entropic forces that work to push against microtubule sliding (Lansky et al., 2015), we speculate that such forces are likely to arise when Ase1 is in a coasting-like mode and the individual crosslinkers are free to diffuse within the confines of the overlap. A more highly tilted braking mode may drive the low microtubule separation observed. This suggests that a coasting-to-braking transition may be more generally applicable to MAP65 family members such as Ase1 and PRC1. The crosslinking motor human kinesin-5/KIF11 drove a lower separation of microtubule pairs when sliding apart antiparallel microtubules compared to crosslinking parallel microtubules, and the separation decreased for faster antiparallel sliding (Meißner et al., 2024). This illustrates low separation/high tilt occurring for higher force generating states of a sliding motor. However for kinesin-14/Ncd, microtubule separation was independent of sliding velocity (Mitra et al., 2020), suggesting that the relationship between separation and force may depend on the crosslinking molecule.

It is possible that the regulation of discrete changes in microtubule spacing could be important in other cellular contexts. For example, during ciliary beating, dynein motors exert force and provide a power stroke which forces the elastic microtubules to bend, likely altering microtubule spacing relative to a relaxed state (King and Sale, 2018). Crosslinking proteins, which include nexin and radial spoke protein complexes that stabilize microtubules against mechanical stress, must compensate for this active force and provide resistance against filament sliding. After the power stroke, the cilium relaxes back to its initial position and relative microtubule arrangement. It is possible that during the transition from power stroke to relaxation mode, the network of crosslinkers subject to bending forces transition between different configurations as the microtubule network undergoes distinct cycles of loading and unloading. Similar principles may be relevant in neuronal microtubule networks as well, where MAPs such as tau and MAP2 establish distinct spacing between adjacent microtubules (Chen et al., 1992). Under mechanical stress this spacing may be modulated due to MAP rearrangement (Chung et al., 2016), and it will be useful to see whether this is a continuous or discrete transition.

These results broaden our understanding of how nanometer-sized proteins that crosslink microtubules collectively give rise to micron-scale mechanical outputs such as resistive force, addressing a challenge in cytoskeletal function. The two-state resistive behavior we identify as controlled by lateral microtubule spacing illustrates the importance of the three-dimensional geometry of microtubule networks in regulating cytoskeletal assemblies. The mechanisms by which diverse crosslinking proteins of differing length compete for the same substrate to drive network mechanics like filament sliding is at present poorly understood. Expanding this work in the future to include additional crosslinkers and microtubule regulators would begin to answer long-standing questions about cytoskeletal organization, not just within the anaphase spindle midzone, but across diverse filament networks in cells.

## METHODS

We performed simulations using the Cytoskeletal Lattice-based Kinetic Simulator (CyLaKS) (Fiorenza et al., 2021). PRC1 molecules and microtubules were explicitly modeled while the kinesin-driven sliding velocity was found by calculating the relationship between sliding velocity and resistive PRC1 force, as discussed below. During each time step a Monte Carlo substep for the kinetic processes determines state change events including PRC1 binding, unbinding, and diffusive hopping. Then a Brownian dynamics (BD) substep computes the motion of the microtubules under force. Here we describe extensions to previous work (Fiorenza et al., 2021).

### PRC1 energy and force

In previous work, crosslinkers were modeled with a linear spring response to compression and extension (Fiorenza et al., 2021), exerting a force on each head *F*_*L*_ = −*k*_s_(*ℓ* − *ℓ*_0_) along the molecular axis, where *ℓ* is the PRC1 head-to-head distance and *ℓ*_0_ the relaxed length. We extended this to include a torque to restore the PRC1 to a preferred angular orientation, motivated by previous work showing a prevalent tilt angle in PRC1 molecules (Subramanian et al., 2010; Kellogg et al., 2016). Structural predictions using AlphaFold also indicate that the microtubule-binding spectrin domain tilts towards the plus end of the microtubules (Fig. S1). We also note that a hinge-like region between the spectrin domain and the coiled-coil domain (Fig. S1) could give rise to a possible asymmetric torsional response of the PRC1 when disturbed from its preferred tilt angle. We use a spring model to express the restoring torque as proportional to the difference between the PRC1 angle *ϕ* and the preferred angle *ϕ*_0_ (measured relative to the normal to the microtubule lateral surface). We implemented a model with directional asymmetry in the restoring torque

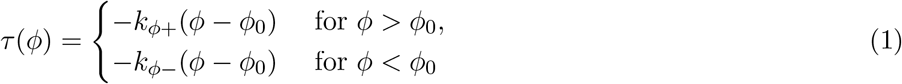

with distinct angular spring constants *k*_*ϕ*+_ and *k*_*ϕ*−_ for the response against tilts toward the positive, respectively negative, angular directions. This choice led to model results more consistent with experimental observations (Fig. S4, S6). This restoring torque induces a force *F*_*T*_ = 2*k*_*ϕ*±_(*ϕ* − *ϕ*_0_)*/ℓ*_0_ on each head normal to the PRC1 molecular axis. Cooperativity in PRC1 binding as previously observed (Subramanian et al., 2010; van den Wildenberg, 2011) is represented by an attractive neighbor-neighbor interaction energy of magnitude *N*_*n*_*ϵ* per PRC1 molecule where *N*_*n*_ is the number of neighboring PRC1 heads bound at adjacent sites along the microtubule. The total potential energy of a crosslinking PRC1 molecule is the sum of the linear spring energy, angular spring energy, and neighbor-neighbor interaction energy

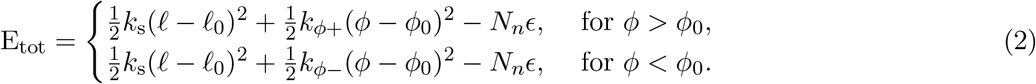

For PRC1 with one head bound, only the neighbor-neighbor interaction energy term is present.

Because PRC1 crosslinkers are passive molecules, their kinetics satisfy detailed balance. We implement this in the rates of binding, unbinding, and diffusion as (Fiorenza et al., 2021),

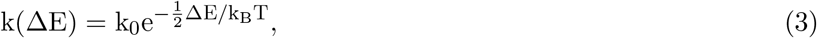

where k(ΔE) is the rate of a state transition, k_0_ is the base rate if no energy change occurs (with value from Table 1), k_B_ is the Boltzmann constant, *T* is room temperature, and ΔE is the change in energy between the current state of the PRC1 molecule and the state after the transition.

### Microtubule force and sliding

Microtubules experience lateral and longitudinal force from PRC1 molecules, kinesin-driven sliding, and steric interactions. We modeled repulsive steric interactions between microtubules as a function of the lateral distance r between their centers using the Weeks-Chandler-Anderson (WCA) potential (Weeks et al., 1971)

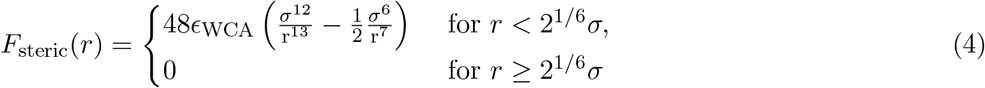

where *ϵ*_WCA_ is the strength of the potential and *σ* the interaction distance.

The surface-attached kinesin motors slide microtubules at a speed that depends on their loading in response to the force exerted by the crosslinking PRC1 molecules. This could be modeled via explicit modeling of the kinesin-microtubule interactions as in previous work (Karan and Chaudhuri, 2023; Arpağ et al., 2014; Palacci et al., 2016), but we prefer to focus the computational model on the crosslinkers, and represent the microtubule as effective boundary conditions for the PRC1 molecules. Because the microtubule dynamics are overdamped, we seek to describe the speed of a microtubule driven by kinesin as a function of the force applied to it by the crosslinking PRC1. In the usual gliding assay paradigm, we would expect in the absence of crosslinking forces for the microtubule to move at approximately the speed *V* at which the unloaded kinesin would walk on a microtubule, because the drag force on the microtubule is small relative to the motor stall force *F*_s_ (Palacci et al., 2016; Leduc et al., 2007; Hunt et al., 1994). When PRC1 crosslinks antiparallel microtubules that are slid apart by kinesin (Fig. 1E), the PRC1 will be stretched and exert forces typically both resisting sliding and pulling the microtubules together. The PRC1 longitudinal force and the (relatively small) drag force on the microtubule must be balanced by the kinesin driving force. From the perspective of a kinesin motor, these forces cause tension in the tether between the kinesin heads and the fixed attachment to the slide, which is oriented against the walking motion of the motor. The walking speed *V* of kinesin-1 is slowed by a resisting force *F*_∥_; we take a widely used piecewise linear force-velocity relation:

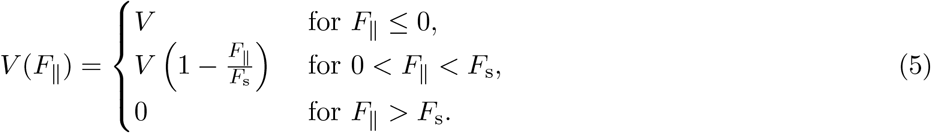

Because *F*_∥_ is the force on a single kinesin, if the total force *F*_*L*_ exerted by the PRC1 on a microtubule is shared equally by *N*_kin_ kinesins, each motor would step at speed *V* (*F*_*L*_*/N*_kin_). This gives a self-consistent deterministic equilibrium with the microtubule moving at velocity *u*_MT_ = *V* (*F*_*L*_*/N*_kin_). However, both experimental and theoretical studies (Arpağ et al., 2019; Driver et al., 2011; Klumpp et al., 2015) of cooperation of multiple kinesin-1 acting on a common cargo provide no reason to expect that the kinesin will organize into such a locked and synchronized quasi-equilibrium.

We thus adopt the more physically plausible framework presented by Karan and Chaudhuri (2023) in which the kinesins are assumed to function independently of each other, apart from the condition that the microtubule is moving at a common velocity *u*_MT_(*t*) relative to the fixed anchorage points of the motors. This work accounts for some cooperativity in the kinesins, treated phenomenologically. Their fit to data suggested a high degree of synchrony between motors, but no discussion is given to counter the experimental assessments of no observable motor cooperativity in experiments (Leduc et al., 2007). We thus assume independent stochastic kinetics for each motor; for simplicity we fix the number of kinesins that can interact with the microtubule at a fixed value *N*_kin_ based on the length of the microtubule. This assumes that as the microtubule moves out of range of one motor, it moves into range of another one; we neglect stochastic fluctuations in the number of kinesin available to bind to the microtubule.

We represent the microtubule-kinesin interactions as occurring in a plane through the microtubule and perpendicular to the slide, assuming the kinesins are all attached to the slide directly below the microtubule. Of course, in reality, there should be an off-axis component of the tether stretch (Palacci et al., 2016), but we exclude this in the model to avoid introducing extra parameters with a limited basis for meaningful quantification (Block et al., 2003). We neglect fluctuations in the distance *s*_⊥_ between the microtubule and the slide to which the kinesin-1 constructs are attached; the fixed value *s*_⊥_ = 25 nm is assumed to arise from balance between tension of the kinesin-1 tails and steric hindrances including antibodies (Kerssemakers et al., 2006). Thus, the state of each of the *N*_kin_ kinesins assumed to be within range of the microtubule can be bound or unbound. When bound, each motor has a signed displacement *s*_∥_ of the kinesin-1 head from its attachment point (with positive values corresponding to the natural walking direction toward the + end). When the kinesin is attached to the microtubule, we model the tensile force between the head and the tail as a function of the end-to-end distance 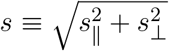 via the cable model

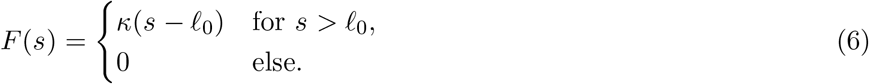

with spring constant *κ* when extended beyond the slack length *ℓ*_0_.

The motor tether force affects kinesin binding kinetics. We denote the force components *F*_∥_ for the longitudinal force and *F*_⊥_ for the transverse force (with *F*_∥_ *>* 0 for forces resisting the walking direction of the motors). The forward hopping rate of bound kinesins is *V* (*F*_∥_)*/δ* with stepsize *δ*. The motor detachment rate is

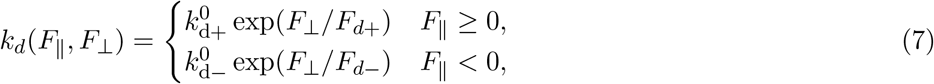

using the experimental data fitting from Andreasson et al. (2015a) but reinterpreted as depending primarily on lateral rather than longitudinal forces, following later work (Pyrpassopoulos et al., 2020; Ma et al., 2023). Unbound mot s attach at a longitudinal displacement randomly distributed according to the Boltzmann distribution 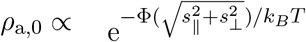 where Φ = ∫ ^*s*^ *F* (*s*) d*s* is the stretching potential energy of the motor tether. Parameters value are given in Table 2.

For a given fixed microtubule speed *u*_MT_, we can calculate the average force a kinesin would exert on the microtubule by computing the stationary distribution of its state via numerical solution of the forward Kolmogorov equation of the stochastic model. This gives the relation between the microtubule velocity *u*_MT_ and *F*_ext_*/N*_kin_, the force exerted on the microtubule per available motor. Further details of this analysis can be found in Kramer (2024). These calculations produce the relations *u*_MT_(*t*) = v_kin_(*F*_ext_(*t*)*/*N_kin_) between the force per kinesin-1 *F*_ext_(*t*)*/N*_kin_ and the microtubule speed *u*_MT_(*t*) for three different values of the unloaded kinesin-1 velocity *V* corresponding to typical values (Schnitzer et al., 2000) for the ATP concentrations used in the experiments in Alfieri et al. (2021) (Fig. 1F). Stall force is held fixed, independent of ATP concentration (Carter and Cross, 2005). Our model gives a substantially lower microtubule velocity under load than the synchronized model discussed above; in particular, the microtubule will stall at substantially lower PRC1 force. This occurs in part because motors are sometimes detached and not applying force, but also significantly because attached motors are often exerting much lower force than they would at mechanical equilibrium.

When PRC1 exerts more force than the average force *F*_max_ the motors are able to apply to keep the microtubule stationary (*u*_MT_ = 0), motors can no longer balance the PRC1 force and the microtubule backslides (against the direction of motor driving). In this case, we assume that the microtubule moves under drag with velocity

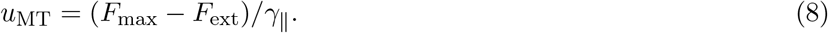

This movement tends to relax the excess PRC1 force rapidly via backsliding, due to the low longitudinal microtubule drag *γ*_∥_. Therefore, we did not implement a detailed model for the kinesin response to the force above stall.

### Time stepping

PRC1 kinetics are simulated with probabilistic sampling of potential events over the time step (Fiorenza et al., 2021), using the kinetic rates of Tables 1,2 modified by the potential energy of the PRC1 molecule as in Eq. (3).

During each BD substep the total force on microtubules from PRC1 molecules and steric interactions between microtubules is calculated. Then the microtubule sliding velocity is calculated in terms of the total longitudinal force 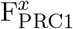 and the number of available kinesins N_kin_ via the force-velocity relation 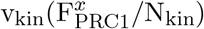(Fig. 1F). The change in longitudinal position *x* and lateral position *y* of each of the microtubules during a time step Δ*t* is calculated using

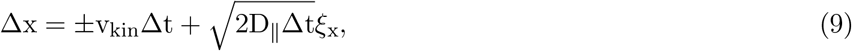

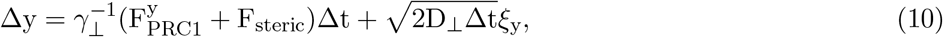

where *γ*_⊥_ is the lateral drag coefficient of the microtubule, *D*_∥_ and *D*_⊥_ are the parallel and perpendicular effective diffusion coefficients of the microtubule, *ξ*_*x*_ and *ξ*_*y*_ are independent standard normal random variables, 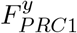 is the lateral component of the force exerted by the crosslinking PRC1 molecules, and *F*_steric_ is the lateral steric force from the other microtubule. The ±*v*_*kin*_ notation indicates that the kinesin-driven microtubule velocity has opposite signs for the two antiparallel microtubules; in our figures we generally have the top microtubule moving to the left.

The maximum velocity v_max_ of the microtubule at zero load corresponds to the escaped velocity of Alfieri et al. (2021). While in principle this should be approximately equal to the unloaded velocity of a single kinesin, the experiments found a broad distribution of escaped velocity. This could arise from motor heterogeneity (Schnitzer et al., 2000; Visscher et al., 1999) and other sources of experimental variability. To represent this experimental variation, in each simulation we independently sample two values of *v*_max_ from a normal distribution with mean and standard deviation of the experimental data of escaped velocity reported in Alfieri et al. (2021, Fig. 2E) for a given ATP concentration, and assign those as the maximum gliding velocity of each of the two antiparallel microtubules. We then obtain the force-velocity relationship of the microtubule by linear interpolation of the coefficients of quadratic fits to the three reference plots in Figure 5B to the sampled values of *v*_max_ in the current simulation.

### Initialization

We initialized simulations with two antiparallel microtubules laterally separated by 30 nm (between surfaces) and longitudinally aligned with an initial overlap length of 5 *μ*m. We determined the number of initially bound PRC1 molecules by sampling from the a Poisson distribution with an expectation of 100 molecules (Table 1). Each PRC1 molecule was attached by crosslinking a pair of directly opposing sites (one on each microtubule) chosen uniformly at random without replacement. We then simulated binding equilibration of immobile overlaps for t_equil_ of 1.5 second, during which sliding was disabled. Sliding then began and the simulations continued until the microtubules were no longer overlapping or until the simulations reached the maximum time t_f_.

For simulations with PRC1 varied between overlap edges and bulk (Fig. 4E-G), the total number of crosslinkers was sampled from the same Poisson distribution used for other simulations. Then the selected number of edge PRC1 molecules was placed by inserting PRC1 starting at each edge and sequentially crosslinking each adjacent pair of opposing sites the specified number of times. The remaining PRC1 molecules were then inserted randomly to the remaining empty pairs of opposing sites as above.

### Data analysis

The average PRC1 occupancy was computed by smoothing the PRC1 data in both time and space. The locations of the PRC1 molecules at a given moment of time *t* were represented by a kernel density estimate using a Gaussian kernel centered at the simulated values over a data collection time step [*t, t*+*t*_*c*_] with a standard deviation of 100 nm for occupancy measures from a single simulation, chosen to be similar to fluorescence imaging experiments. When averaging many simulations, we used a standard deviation of 30 nm (shown in Fig. 2E-I, 4F,G). The resulting occupancy function was normalized so that a value of 1 corresponds to full occupancy of available PRC1 binding sites. The edge region of the microtubule overlaps was defined as the 240 nm from the end of the microtubule defining the overlap edge. This was chosen as a typical width for the PRC1 clusters on overlap edges in our simulations and in experiments (Alfieri et al., 2021).

### Parameters

The values of the parameters used in the computational model and the kinesin-microtubule interaction model are listed in Tables 1 and 2, respectively. When those values are taken from or motivated by the literature, the source is indicated. Some parameters were chosen by computational parameter searches to fit the experimental data; these are indicated as fit and discussed below.

The longitudinal diffusion of microtubules in a gliding assay with a single attached kinesin has been estimated from experiment (Palacci et al., 2016) to be 1400nm^2^*/*sec at saturating ATP concentration, an order of magnitude smaller than the longitudinal diffusion coefficient of freely diffusing microtubules. This microtubule diffusion appears to primarily come from the kinesin tethering the microtubules to the surface, and to be insensitive to the microtubule length (Imafuku et al., 2008). The diffusion coefficient has been found to decrease somewhat at higher kinesin density, but our linear density of available kinesin 6 *μ*m^−1^ is lower than the densities previously studied (Palacci et al., 2016). We thus take 1400 nm^2^*/*sec as a baseline microtubule diffusion coefficient for saturating ATP. Assuming the longitudinal diffusivity to scale linearly with the motor velocity, as in standard motor stepping models, we could choose the longitudinal diffusivity of the microtubules to be adjusted downward from the ATP-saturated value as

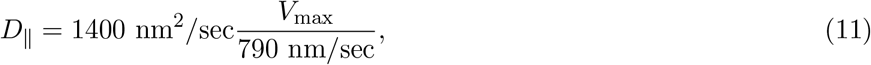

with the denominator the kinesin-1 velocity at saturating ATP (Andreasson et al., 2015b). Note we adjust the microtubule diffusivity only based on the maximum microtubule sliding velocity, and not on the velocity under load. This model neglects the PRC1 crosslinkers, which presumably would further reduce the microtubule diffusion. We found that reducing the longitudinal diffusivity formula in Eq. (11) by a factor of 10 gave good agreement between the simulation results and the experimental data. We are unaware of experimental measurements or theory for lateral diffusion of gliding microtubules, so we chose *D*_⊥_ = *D*_∥_*/*2 as for freely diffusing microtubules. While these are rough estimates, because the microtubule diffusivity is at least an order of magnitude smaller than the diffusivity of doubly-bound PRC1, the precise parameterization is not expected to play a significant role.

The coupling of the microtubule to surface-attached kinesin and PRC1 would be expected to increase the effective drag coefficient from that of a freely floating microtubule, since they slow the longitudinal diffusion. But a precise determination for the drag increase does not appear available. Indeed, in actively driven systems, the Einstein relation along with the classical fluctuation-dissipation relationships only applies in limited settings with a different proportionality that may not be strongly dependent on temperature (Caprini et al., 2021; Fodor et al., 2016). Fortunately, a consideration of time scales implies that the simulation should not be so sensitive to the precise drag values, as we now explain. Using the standard fluid mechanics formula for the drag of a cylinder near a planar surface in a fluid with the viscosity of water(Howard, 2001) gives a drag coefficient that would result in a very rapid longitudinal backsliding of the microtubule under excess PRC1 force, requiring a much reduced time step to resolve. We therefore use instead the Einstein relation to specify *γ*_∥_ = *k*_*B*_*T/D*_∥_ and *γ*_⊥_ = *k*_*B*_*T/D*_⊥_, which we rather expect to be an overestimate. But the drag values obtained in this ad hoc way are suitable for our computational model. With this enhanced microtubule drag, the longitudinal microtubule dynamics still respond rapidly over a couple of time steps to relax excess PRC1 force over the maximum kinesin force without requiring a reduction in time step. The lateral dynamics of separation between the microtubules have a time scale *γ*_⊥_*/*(*N*_2_*k*_*s*_) ∼ 0.01 sec, where *N*_2_ is the number of doubly bound PRC1. This is fast enough to respond to the relatively slow variation in lateral force. For the purposes of our simulations, the microtubule drag coefficient parameters simply need to be small enough for the microtubule to rapidly restore force balance.

The steric interaction distance *σ* for the microtubules was chosen to be 45 nm, causing microtubules to repel each other when closer than r = 2^1*/*6^*σ* = 50.5 nm. This was increased beyond the 25-nm microtubule diameter to model, for example, steric interactions between microtubule C-terminal tails and/or electrostatic repulsion between microtubule surfaces. If *σ* is set to 25 nm microtubules in overlaps become unphysically close while in the braking state, with edge to edge separation of ≈ 5 nm (Fig. S7).

The linear spring constant for PRC1 k_s_ was increased from previous estimates (Wierenga and ten Wolde, 2020) by an order of magnitude to 2 pN/nm to enable sufficient braking. Without this increase, overlaps rarely transitioned to braking (Fig. S8).

The attachment rate of the second PRC1 head is estimated using the procedure from Fiorenza et al. (2021) in which the free head is assumed to explore a half-sphere of radius 32 nm, and thus have an effective concentration of 24700 nM.

The enhanced detachment rate 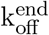 of the PRC1 heads at the microtubule ends decreases the rate at which PRC1 builds up on the edges of overlaps and was chosen so that peaks formed in the simulations at a rate similar to what was observed in experiments (Alfieri et al., 2021) (Fig. S9). If 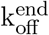 is set too low, overlaps rarely stay in the coasting mode (Fig. S9). With both PRC1’s angular tilt preference and increased end unbinding rate turned off, overlaps displayed braking/coasting behavior, however large PRC1 occupancy peaks were always present at overlap edges even in the coasting mode, at odds with experiment (Fig. S10) (Alfieri et al., 2021).

While we are unaware of any experimentally calibrated descriptions of such behavior, we found that including an explicit asymmetric torsional response (1) to PRC1 rotation relative to the microtubules increased the likelihood of overlaps transitioning to the braking state by preventing PRC1 molecules from being tilted against sliding (Fig. 4,S4). We set the angular spring constant against negative rotation, k_*ϕ*−_, so that overlaps transitioned to braking when moderate PRC1 clusters formed on the edge of overlaps. If k_*ϕ*−_ is too high, all overlaps quickly transition to the braking state. If k_*ϕ*−_ is too low, overlaps rarely transition to braking even after PRC1 clusters at the edge. With a symmetric response, k_*ϕ*+_ = k_*ϕ*−_, the restoring torque from PRC1 prevented overlaps from staying in the braking state (Fig. S6). We therefore set the angular spring constant against positive rotation, k_*ϕ*+_, lower than k_*ϕ*−_.

The number of kinesin driving a microtubule in the experiments of Alfieri et al. (2021) was inferred by finding the critical kinesin concentration to get only 1 kinesin attached to the microtubule using previously reported methods (Howard et al., 1989), and then rescaling to the bulk kinesin concentration used in the experiments to study the braking behavior of PRC1. A microtubule was deemed to have exactly 1 kinesin attached when the microtubule exhibited directional transport without bending at low bulk kinesin concentration. This procedure suggested a linear density of roughly 5 kinesin per micrometer. We chose a somewhat higher density of kinesin that are available to the microtubule, since the calculations in Kramer (2024) indicate that the available kinesin remains attached to the microtubule about 70% of the time.

The means and standard deviation for the escaped velocities *V*_max_ are taken at the three ATP concentrations 10 *μM*, 100 *μ*M, and 500 *μ*M as 127 ± 37 nm/sec, 254 ± 68 nm/sec, and 410 ± 98 nm/sec respectively (Fig. 5B). Most simulations correspond to the 10 *μ*M model. The higher sliding speeds are used in simulations of Fig. 5B-C.

The rest length for kinesin is estimated based on the truncated form K439 used in the experiments of Alfieri et al. (2021), and measurements of similar constructs in Kerssemakers et al. (2006); Woll et al. (2018).

## Supplemental information index

Figures S1-S9 and their legends in a PDF.

## Acknowledgments

This work was funded by the NSF via grant DMS 2153399 to MB, DMS 2153374 to PRK and SForth, and via NIH/NIGMS grant R01GM149782 to SForth.

## Author contributions

Conceptualization, ET, SForth, PRK, MB; methodology, DS, SFiorenza, PRK, MB; investigation, DS, SFiorenza, ET, SForth, PRK, MB; writing, original draft, DS, SFiorenza, SForth, PRK, MB; writing, review & editing, DS, ET, SForth, PRK, MB; funding acquisition, SForth, PRK, MB; resources, SForth, PRK, MB; supervision, SForth, PRK, MB.

## Declaration of generative AI and AI-assisted technologies

During the preparation of this work, the authors used no AI technologies.

