## Supplemental figures for "PRC1 resists microtubule sliding in two distinct resistive modes due to variations in the separation between overlapping microtubules"

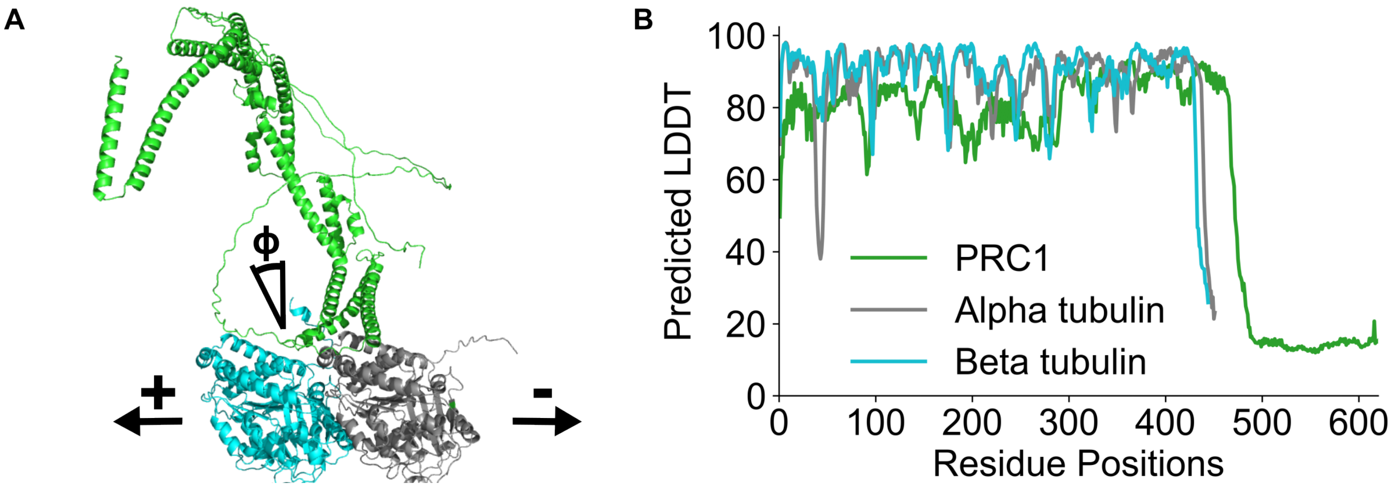


FIG. S1. **AlphaFold predictions suggest a structural tilt of PRC1 towards the plus end of the bound microtubule.** (A) AlphaFold predicted structure of PRC1 attached to a tubulin dimer with the microtubule plus and minus end directions shown. PRC1 is predicted to tilt toward the plus end of the microtubule. (B) The predicted local distance difference test (LDDT) score is shown for the AlphaFold model. The low confidence region for residues with positions >467 correspond to the disordered C-terminal domain.

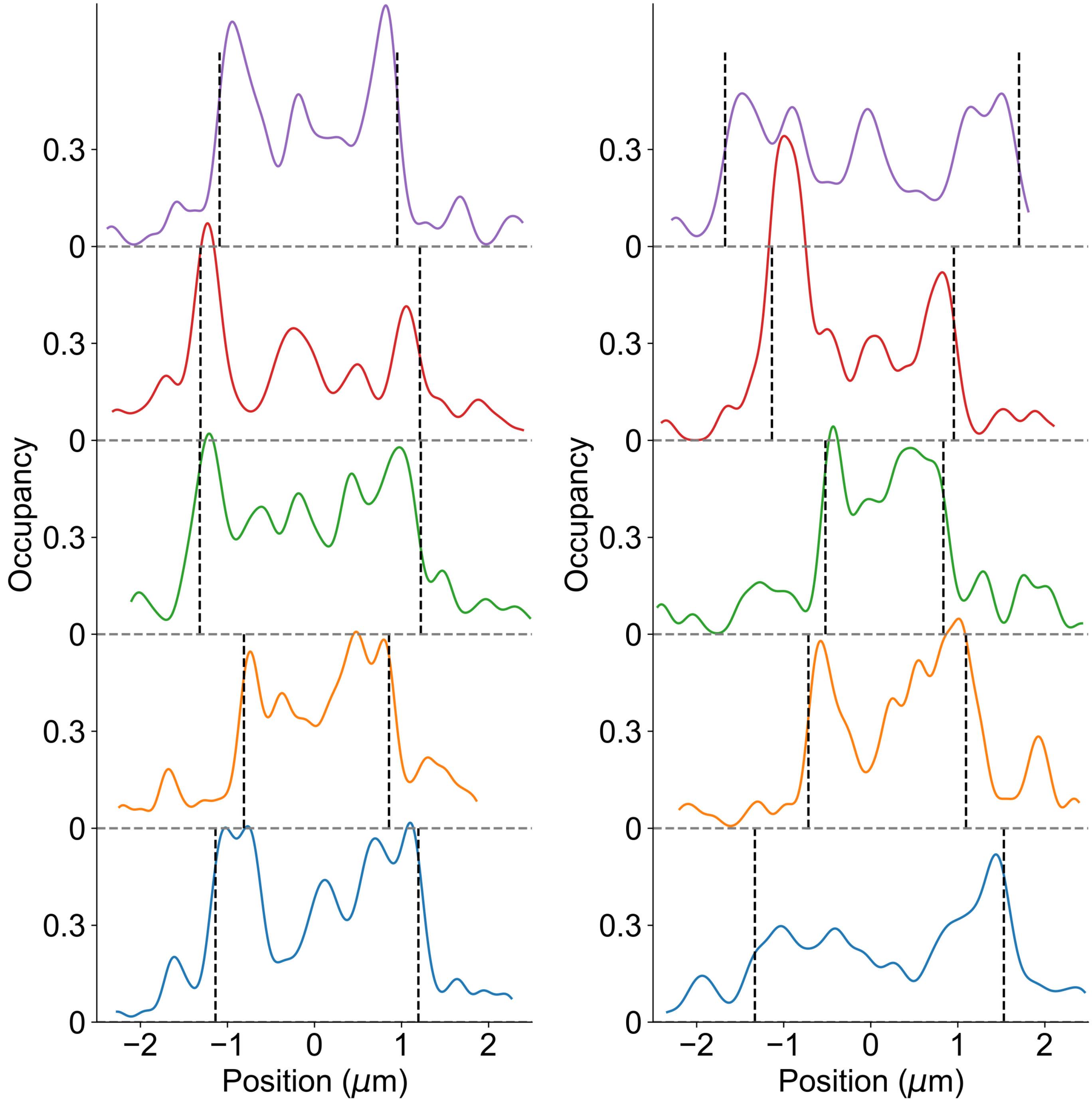


FIG. S2. **PRC1 clusters at overlap edges during transitions to the braking state.**
Each curve shows the PRC1 occupancy for a different overlap averaged over the data collection frame that the overlap transitioned from a separation of greater than 25 nm (coasting) to a separation of less than 25 nm (braking).

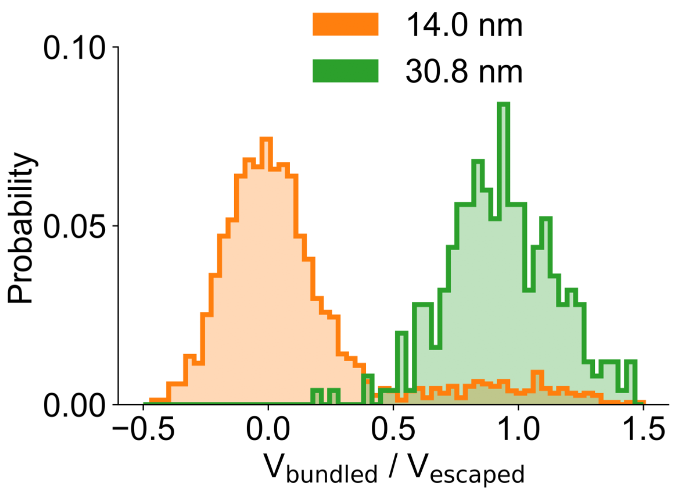


FIG. S3. **Overlaps with separation fixed at 30.8 nm maintained coasting sliding speeds, while overlaps with separation fixed at 14.0 nm maintained braking speeds.**
Distribution of normalized velocity for overlaps with the separation fixed at 30.8 nm and with the separation fixed at 14.0 nm. Data points are each the average over 1 second of the simulation. Each separation has data from 10 simulations. Note: The high normalized velocity data when the separation was fixed at 14.0 nm is from when the overlaps are close to sliding apart.

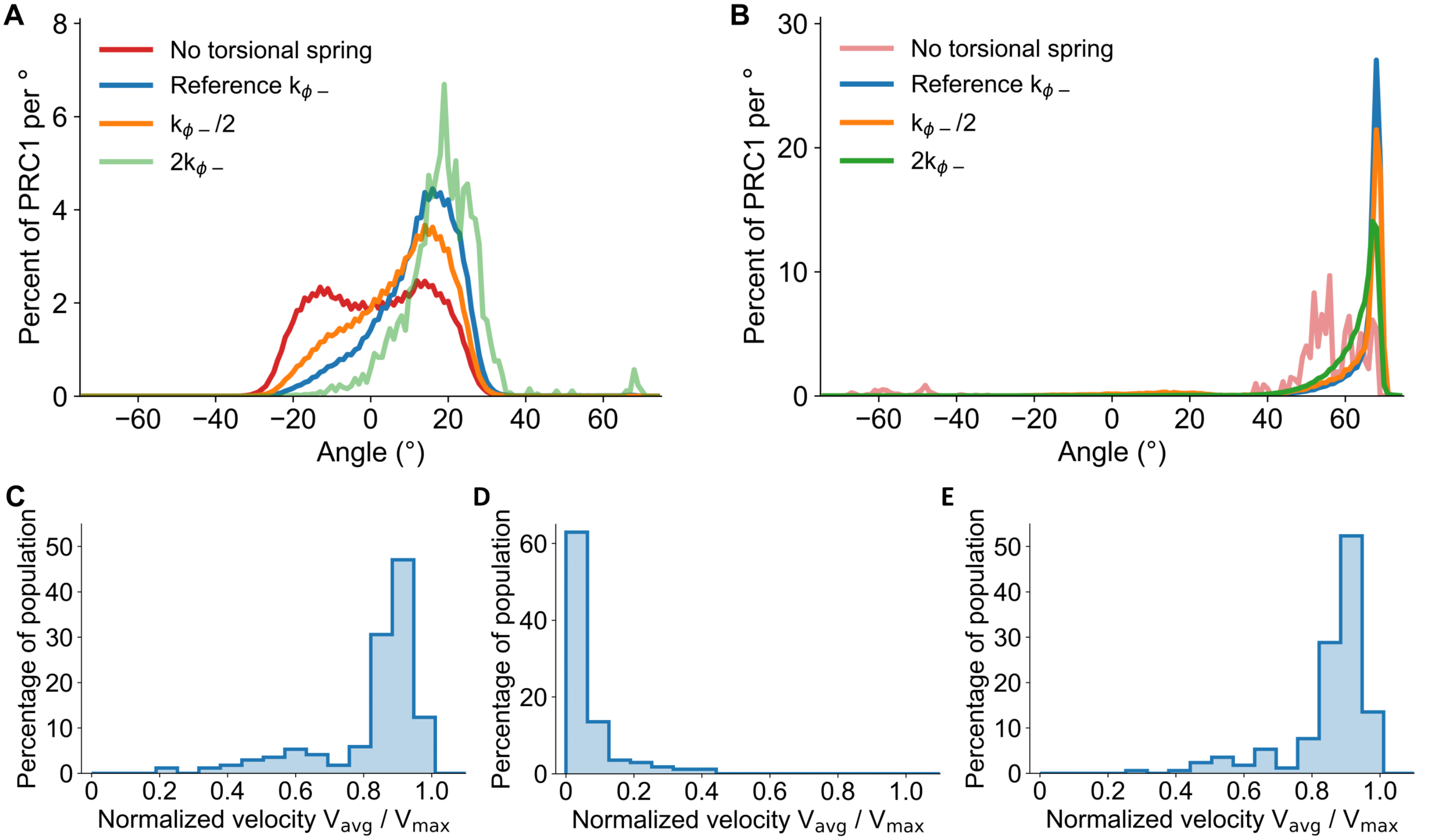


FIG. S4. **Greater resistance to negative tilting promotes PRC1 alignment in the sliding direction and braking.** (A-B) Distribution of PRC1 tilt angle in the coasting state (A) and braking state (B) for simulations with varying torsional spring constant against negative tilt $k_{\phi-}$ (Table 1). (A) For larger $k_{\phi-}$ increases, more PRC1 molecules have positive tilt during coasting. (B) During braking PRC1 molecules are consistently tilted with sliding (C-E) Normalized sliding velocity distribution for simulations with $k_{\phi-}$ reduced to half the reference value (C), increased to twice the reference value (D), or with no torsional force ($k_{\phi-}$=$k_{\phi+}$=0, E). Data from 100 simulations for each parameter set.

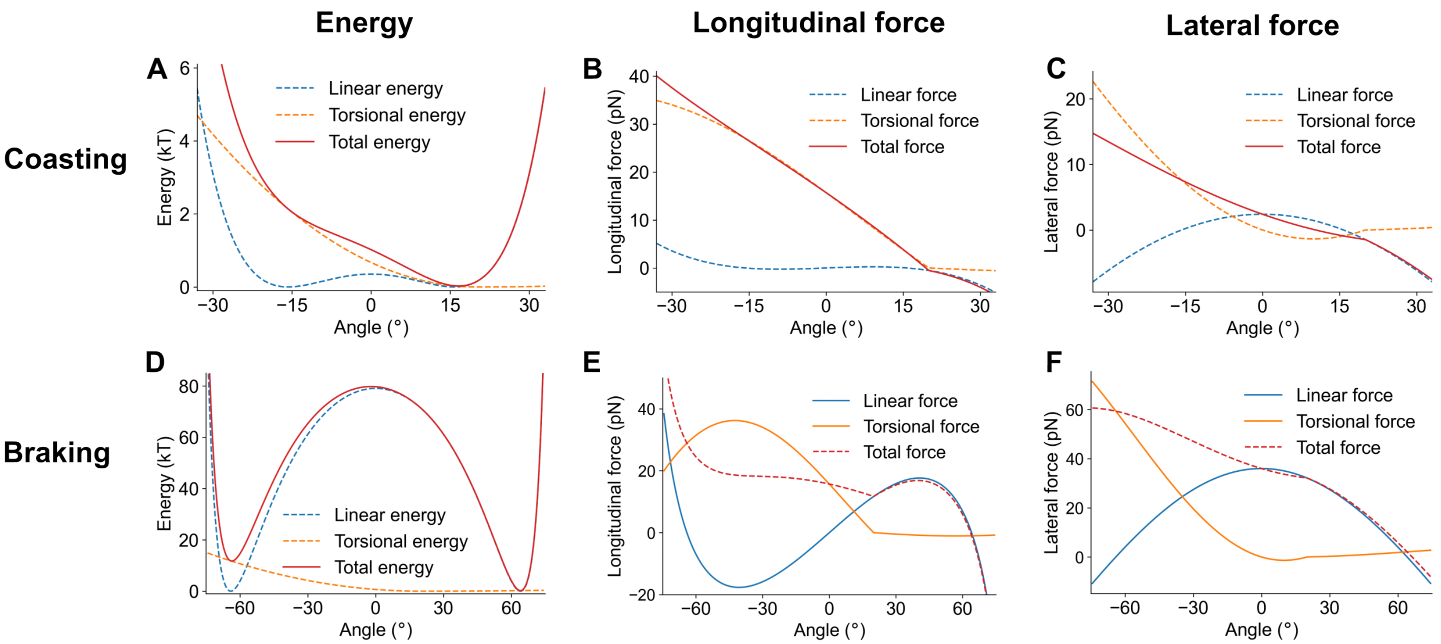


FIG. S5. **The torsional and linear forces both contribute to the longitudinal and lateral forces produced by PRC1** (A) The linear, torsional, and total energy of an individual PRC1 molecule calculated at coasting separation (30.8 nm). (B-C) The linear contribution, torsional contribution, and total longitudinal(B)/lateral(C) force for a PRC1 molecule calculated at coasting separation. (D) The linear, torsional, and total energy of an individual PRC1 molecule calculated at braking separation (14.0 nm). (E-F) The linear contribution, torsional contribution, and total longitudinal(E)/lateral(F) force for a PRC1 molecule calculated at braking separation.

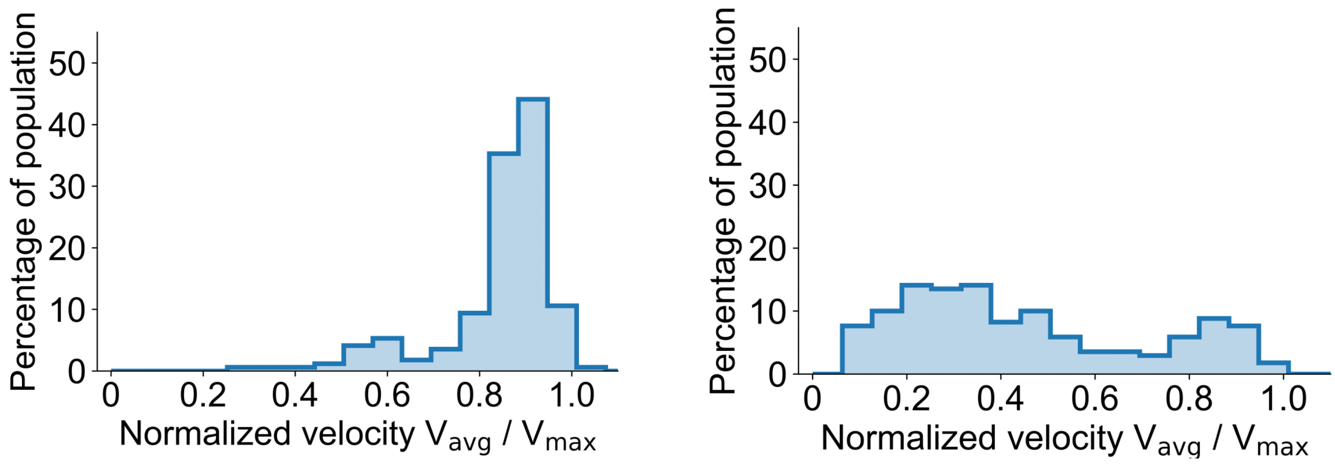


FIG. S6. **Greater resistance to positive tilting promotes coasting.**
Normalized sliding velocity distribution for overlaps with $k_{\phi+}$ increased to the same value as $k_{\phi-}$ (A) or decreased to 0 (B).

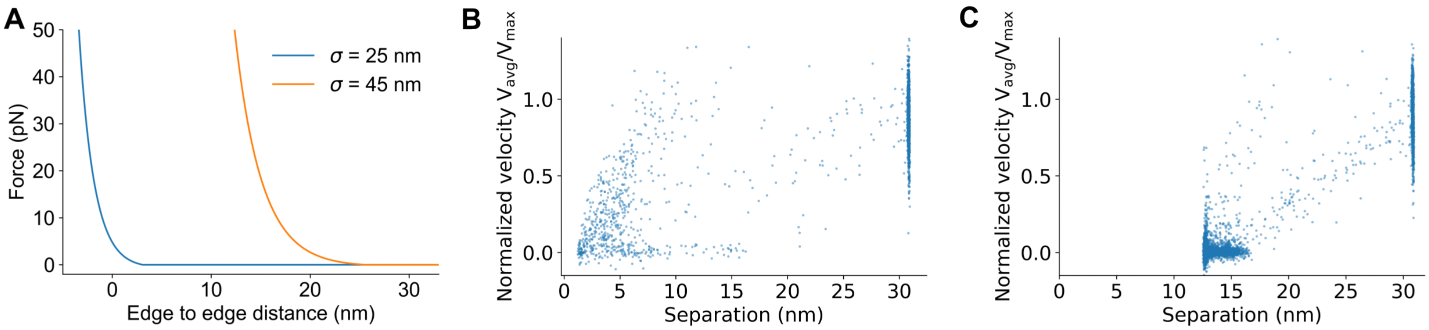


FIG. S7. **The WCA potential interaction distance tunes microtubule lateral separation.**
(A) The force from the WCA potential as a function of the microtubule separation for an interaction scale equal to the microtubule diameter ($\sigma = 25$ nm) and for our chosen interaction scale ($\sigma = 45$ nm). (B-C) Normalized overlap sliding velocity versus microtubule pair lateral separation $\sigma = 25$ nm (B) and $\sigma = 45$ nm (C). Each data point is the average from one data collection time interval, collected from 100 simulations.

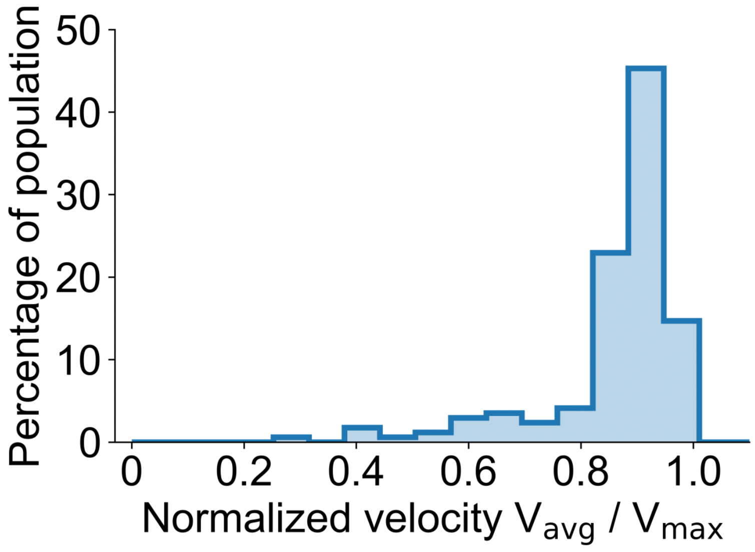


FIG. S8. **Decreased PRC1 linear spring constant favors coasting.**
Normalized sliding velocity distribution for PRC1 linear spring constant lowered to half the reference value. Data from 100 simulations.

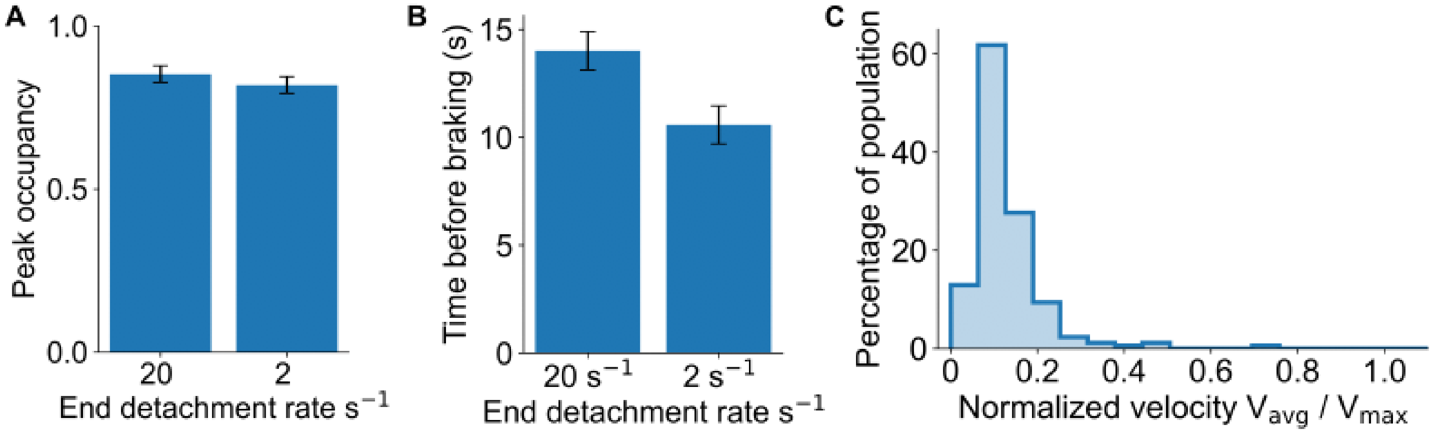


FIG. S9. **Increased PRC1 lifetime at overlap edges promotes braking.**
(A) Peak occupancy at the time of transition from coasting to braking for the reference end unbinding rate $k_{off}^{end} = 20s^{-1}$ and a factor of ten lower end unbinding rate $k_{off}^{end} = 2s^{-1}$. (B) The time between the start of sliding to the transition from coasting to braking for the reference and decreased end unbinding rate. The time is decreased when PRC1 molecules have a longer lifetime at the overlap edge. (C) Normalized velocity distribution for overlaps with decreased end unbinding rate $k_{off}^{end} = 2s^{-1}$. Data from 100 simulations for each parameter set.

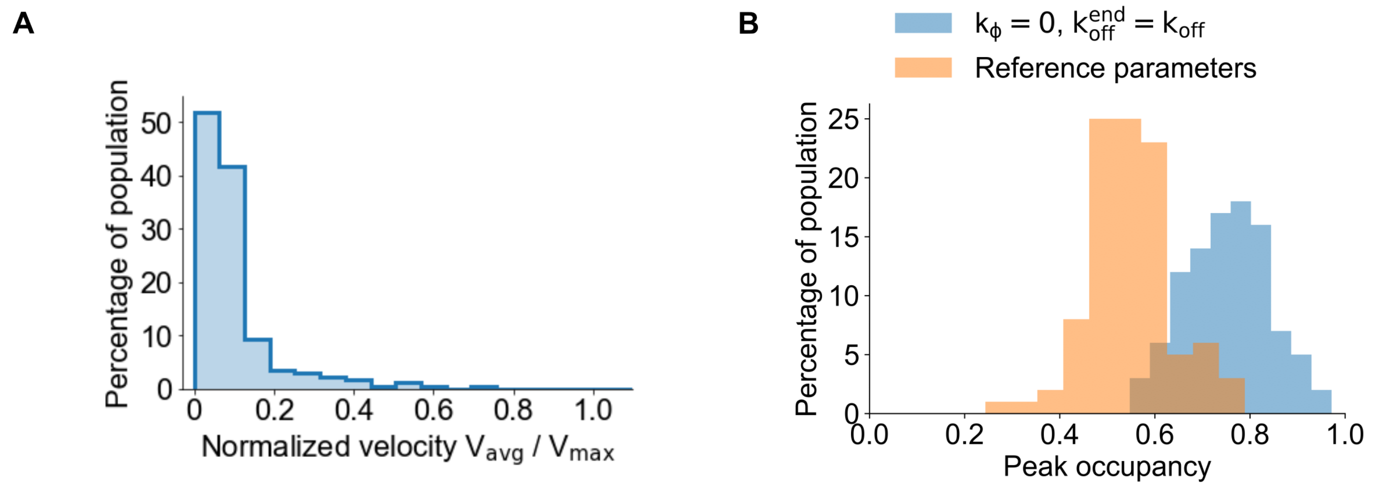


FIG. S10. **Braking can occur with an altered parameter set but the PRC1 distribution then differs from experimental observations.**
(A) Normalized velocity distribution for overlaps without enhanced PRC1 end unbinding and without a PRC1 restoring torque. Overlaps frequently show braking. (B) The distribution of the average overlap edge occupancy while coasting for simulations with the reference parameters and for simulations without enhanced end unbinding or a restoring torque. For these parameters, most overlaps in the coasting state show PRC1 clusters near overlap edges, in contrast to the experimental data. Data from 100 simulations for each parameter set.
